# Functional genomics reveals extensive diversity in *Staphylococcus epidermidis* restriction modification systems compared to *Staphylococcus aureus*

**DOI:** 10.1101/644856

**Authors:** Jean YH Lee, Glen P Carter, Sacha J Pidot, Romain Guérillot, Torsten Seemann, Anders Gonçalves da Silva, Timothy J Foster, Benjamin P Howden, Timothy P Stinear, Ian R Monk

## Abstract

*Staphylococcus epidermidis* is a significant opportunistic pathogen of humans. Molecular studies in this species have been hampered by the presence of restriction-modification (RM) systems that limit introduction of foreign DNA. Here we establish the complete genomes and methylomes for seven clinically significant, genetically diverse *S. epidermidis* isolates and perform the first systematic genomic analyses of the type I RM systems within both *S. epidermidis* and *Staphylococcus aureus*. Our analyses revealed marked differences in the gene arrangement, chromosomal location and movement of type I RM systems between the two species. Unlike *S. aureus*, *S. epidermidis* type I RM systems demonstrate extensive diversity even within a single genetic lineage. This is contrary to current assumptions and has important implications for approaching the genetic manipulation of *S. epidermidis*. Using *Escherichia coli* plasmid artificial modification (PAM) to express *S. epidermidis hsdMS*, we readily overcame restriction barriers in *S. epidermidis*, and achieved transformation efficiencies equivalent to those of modification deficient mutants. With these functional experiments we demonstrate how genomic data can be used to predict both the functionality of type I RM systems and the potential for a strain to be transformation proficient. We outline an efficient approach for the genetic manipulation of *S. epidermidis* from diverse genetic backgrounds, including those that have hitherto been intractable. Additionally, we identified *S. epidermidis* BPH0736, a naturally restriction defective, clinically significant, multidrug-resistant ST2 isolate as an ideal candidate for molecular studies.

**Importance:** *Staphylococcus epidermidis* is a major cause of hospital-acquired infections, especially those related to implanted medical devices. Understanding how *S. epidermidis* causes disease and devising ways to combat these infections has been hindered by an inability to genetically manipulate “hospital-adapted” strains that cause clinical disease. Here we provide the first comprehensive analyses of the mechanisms whereby *S. epidermidis* resists the uptake of foreign DNA and demonstrate that these are distinct from those described for *S. aureus.* Until now it had been assumed that these are the same. Using these insights, we demonstrate an efficient approach for the genetic manipulation of *S. epidermidis* to enable the study of clinically relevant isolates for the first time.

## Introduction

*Staphylococcus epidermidis* is a ubiquitous coloniser of human skin (1). Invasive medical procedures, specifically insertion of prosthetic devices on which the bacteria can form a biofilm, enables evasion of both antibiotics and the host immune system and has contributed to its rise as a significant nosocomial pathogen. A leading cause of surgical site and central-line-associated bloodstream infections (2), *S. epidermidis* poses a major economic burden (3). In the hospital environment two multi-locus sequence types (MLSTs), ST2 and ST23, account for most clinical disease (4, 5). Three hospital adapted clones (two ST2, and one ST23) were recently demonstrated to be globally disseminated and have evolved to become nearly untreatable though the acquisition of multiple antibiotic resistance determinants and resistance conferring mutations (5). Knowledge of the molecular genetics, pathogenesis and treatment of *S. epidermidis* has been limited by barriers preventing the genetic manipulation of clinically relevant isolates and the assumption that *S. epidermidis* is similar to *S. aureus*.

Restriction-modification (RM) systems have evolved as a form of bacterial immunity that degrades incoming DNA from foreign donors such as bacteriophage (6). Type I and IV RM systems are a significant barrier to genetic manipulation of staphylococci. Type I RM systems are comprised of three host specificity of DNA (*hsd*) genes that encode (i) a specificity protein (HsdS), (ii) modification proteins (HsdM) and (iii) a restriction endonuclease (HsdR). Together these function as a single protein complex in which HsdS determines the DNA target recognition motif (TRM) in which adenine residues are methylated by HsdM, while HsdR cleaves unmodified and non-self-modified DNA (7, 8). Type IV RM systems consist of a single restriction endonuclease that cleaves DNA with inappropriate cytosine methylation (8).

Plasmid artificial modification (PAM) is a method to overcome the barrier imposed by RM systems, where plasmid DNA is passaged through a cytosine methylation deficient *Escherichia coli* host (DC10B) that has been engineered to heterologously express the *hsdMS* system of a staphylococcal strain to be transformed. Plasmid DNA extracted from this *E. coli* host mimics the DNA methylation profile of the target strain, thus enabling introduction of plasmid DNA and subsequent genetic manipulation (9).

Type I RM systems of staphylococci are best understood in *S. aureus*. The distribution of *hsdS* alleles corresponds with clonal complex (CCs) for the 10 dominant *S. aureus* lineages (10). Far less is known about the type I RM systems in *S. epidermidis*. A recent study suggested that *S. epidermidis* type I RM systems adhered to lineage specific groupings like *S. aureus*. However, this inference was based on only four new *S. epidermidis* methylomes (11) plus the one methylome that was already characterised; the ST2 reference genome of strain BPH0662 (12).

Here, we present the first systematic genomic analyses of the type I RM systems in *S. aureus* and *S. epidermidis* and demonstrate how this data can be used to predict functionality of type I RM systems and transformational competence of strains. We show that PAM is a highly efficient method to enable genetic manipulation of *S. epidermidis*, particularly hospital-adapted isolates that possess multiple functional type I RM systems.

## Results and Discussion

### *S. aureus* type I RM systems are lineage specific

We began this study by testing the notion that *S. aureus* type I RM systems are lineage specific. We compared 128 publicly available finished *S. aureus* genome sequences (Table S1A) and confirmed the chromosomal location and structure for type I RM systems in *S. aureus* is highly conserved. One hundred and ten genomes had a single *hsdR* and two copies of *hsdMS* with the first in forward orientation located in the alpha pathogenicity island and the second in the opposite orientation, located within the beta pathogenicity island (10, 13) (Figure 1A). The remaining 18 strains possessed *hsdR* and a single copy of forward oriented *hsdMS* in the alpha pathogenicity island. Five of the 128 strains possessed a third type I RM system at a non-mobile chromosomal location downstream of *lacA*. Twenty-three strains carried an additional type I RM system on a mobile genetic element, 22 were on staphylococcal cassette chromosome (SCC) elements (Figures 2 & S1) and one type I RM system on a plasmid (strain HUV05). Single variants of both HsdR (NCBI protein accession WP_000331347.1; n = 127) and HsdM (WP_000028628.1; n = 222) were demonstrated for the type I RM systems situated in non-mobile chromosomal locations, indicating stable vertical inheritance. Interruptions in *hsdR* and *hsdMS* due to horizontal gene transfer were rarely seen in *S. aureus.* A solitary example of *hsdS* truncation due to insertion of a bacteriophage was noted in Sa17_S6. Most changes were due to SNPs leading to amino acid substitutions (N = 35) or nonsense mutations (N = 31) in *hsdS* (Figure 2). See Table 1 for a comparison of *S. aureus* and *S. epidermidis* type I RM systems.

**Figure 1.**
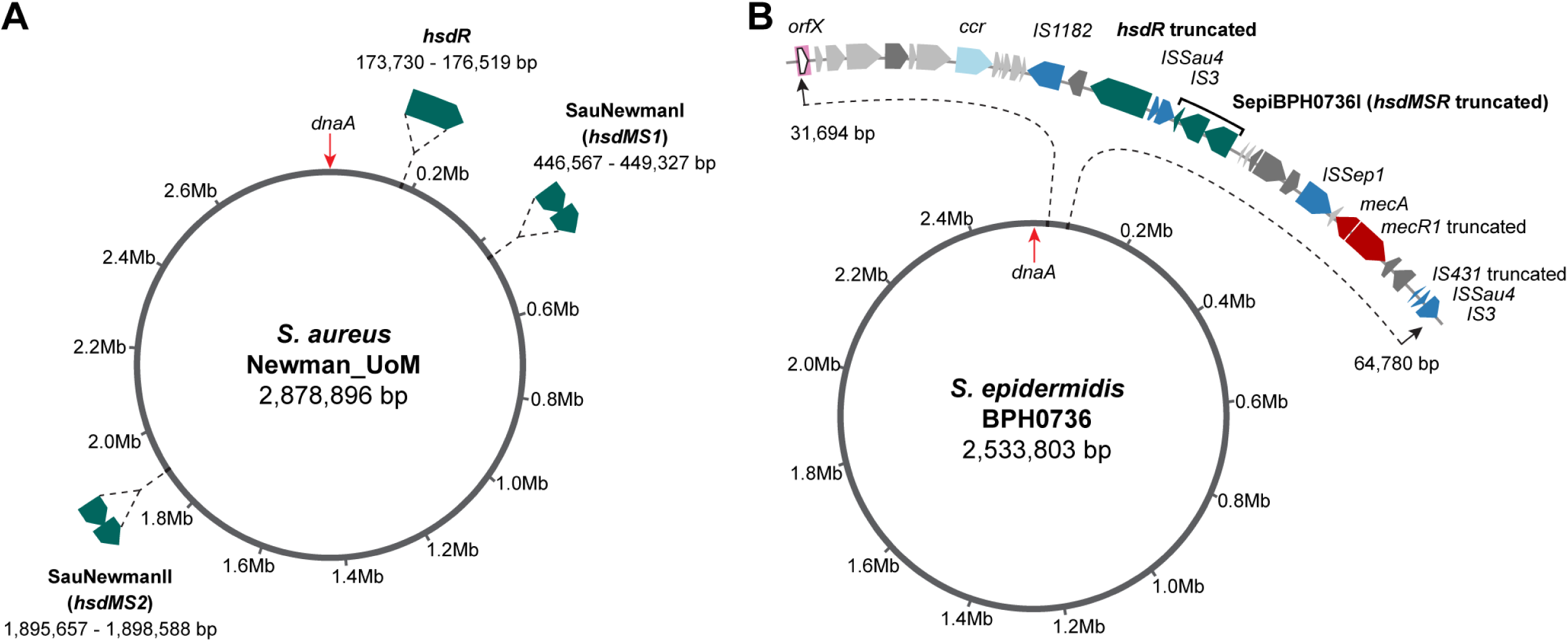
Comparison of the structure and chromosomal location of *S. aureus* and *S. epidermidis* type I restriction modification systems. A. *S. aureus* Newman_UoM (29, 32). B. *S. epidermidis* BPH0736. For consistency, the chromosome was orientated forwards starting at *dnaA* and native type I RM systems were sequentially numbered, with any imported systems numbered thereafter.

**Figure 2.**
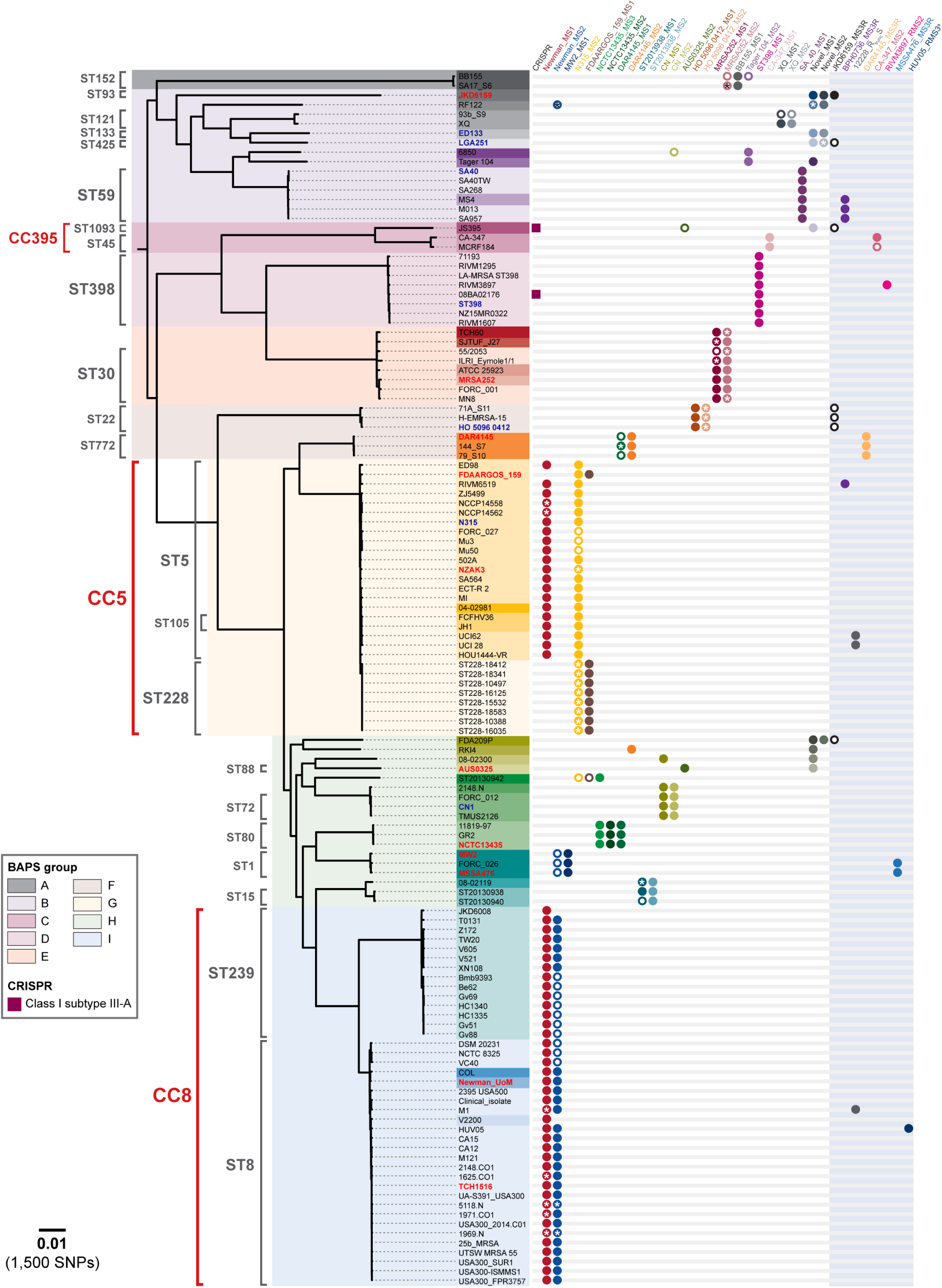
*S. aureus* native type I restriction modification systems are lineage specific. Maximum-likelihood core-SNP based phylogeny of 128 closed *S. aureus* genomes originating from 40 STs, using Newman_UoM as the reference genome. Overlaid are the results of in silico multi-locus sequence type (MLST), clonal cluster (CC), bayesian analysis of population structure (BAPS), presence of CRISPR-Cas systems and type I restriction modification system HsdS variants. Bold red font indicates isolates with PacBio characterised methylomes; Bold blue font indicates isolates with methylomes determined by DNA cleavage with purified enzyme. Boxes around strain names are coloured according ST type. Open circle = amino acid substitutions present in HsdS. *indicates truncated HsdS subunit. Scale bar indicates number of nucleotide substitutions per site (bold) with an approximation of SNP rate (in parentheses).

**Table 1.** Comparison of S. aureus and S. epidermidis type I restriction modification systems.

| *Staphylococcus aureus* | *Staphylococcus epidermidis* |
| --- | --- |
| Organised as a single *hsdR* gene separated from one, two or three distant *hsdMS* gene pairs | Organised as complete three gene operon (*hsdRMS* or *hsdMSR*) |
| Conserved, stable chromosomal location for each gene | Close proximity with *ccr* genes integrated at *orfX*, located in highly plastic region of genome |
| Most have two type I RM systems | Most have a single type I RM system |
| All have at least one type I RM system | Many (38.1%) have no type I RM system |
| Up to three functional type I RM systems per isolate | Up to three functional type I RM systems per isolate |
| 99.7% amino acid pairwise identity for all native HsdR | At least five identified variants of HsdR |
| 99.3% amino acid pairwise identity for all native HsdM | At least six identified variants of HsdM |
| At least 48 different variants of HsdS (eight likely imported from coagulase negative staphylococci) | At least 31 different variants of HsdS |
| Relative conservation of HsdS present within ST groups | No clear conservation of HsdS according to ST group |
| Conservation of HsdM provides redundancy, enabling interaction with multiple different HsdS in *S. aureus* | HsdS are only be capable of interacting with their specific paired HsdM, therefore not all orphan *hsdS* will be functional |
| Complete three gene *hsdRMS/MSR* type I systems carried on SCC elements do not adhere to lineage specificity are likely imported from coagulase negative staphylococci |  |

We next established a high-resolution phylogeny using 144,727 core genome single nucleotide polymorphisms (SNPs) for the 128 *S. aureus* genomes covering 40 STs (Figure 2). The occurrence of *hsdMS* genes for each genome was mapped across the phylogeny. A total of 48 HsdS subunits were identified with associated TRMs (Table 2, Table S2A). Although the same *hsdMS* genes were present in genetically distinct lineages, the combination of *hsdMS* genes were conserved within each lineage (Figure 2). For example, the same two HsdMS were present in ST250 and ST254 which are single locus variants of ST8. A notable exception to the lineage specificity were the type I RM systems carried on SCC elements (Figure S1B) that may have been acquired from coagulase negative staphylococci (CoNS). The complete *S. epidermidis* BPH0736 *hsdMSR* (non-disrupted, identical sequence) was observed in four *S. aureus* strains from three different STs (ST5, ST59 and ST338) suggesting gene transfer between the species (Figure S1).

**Table 2.**
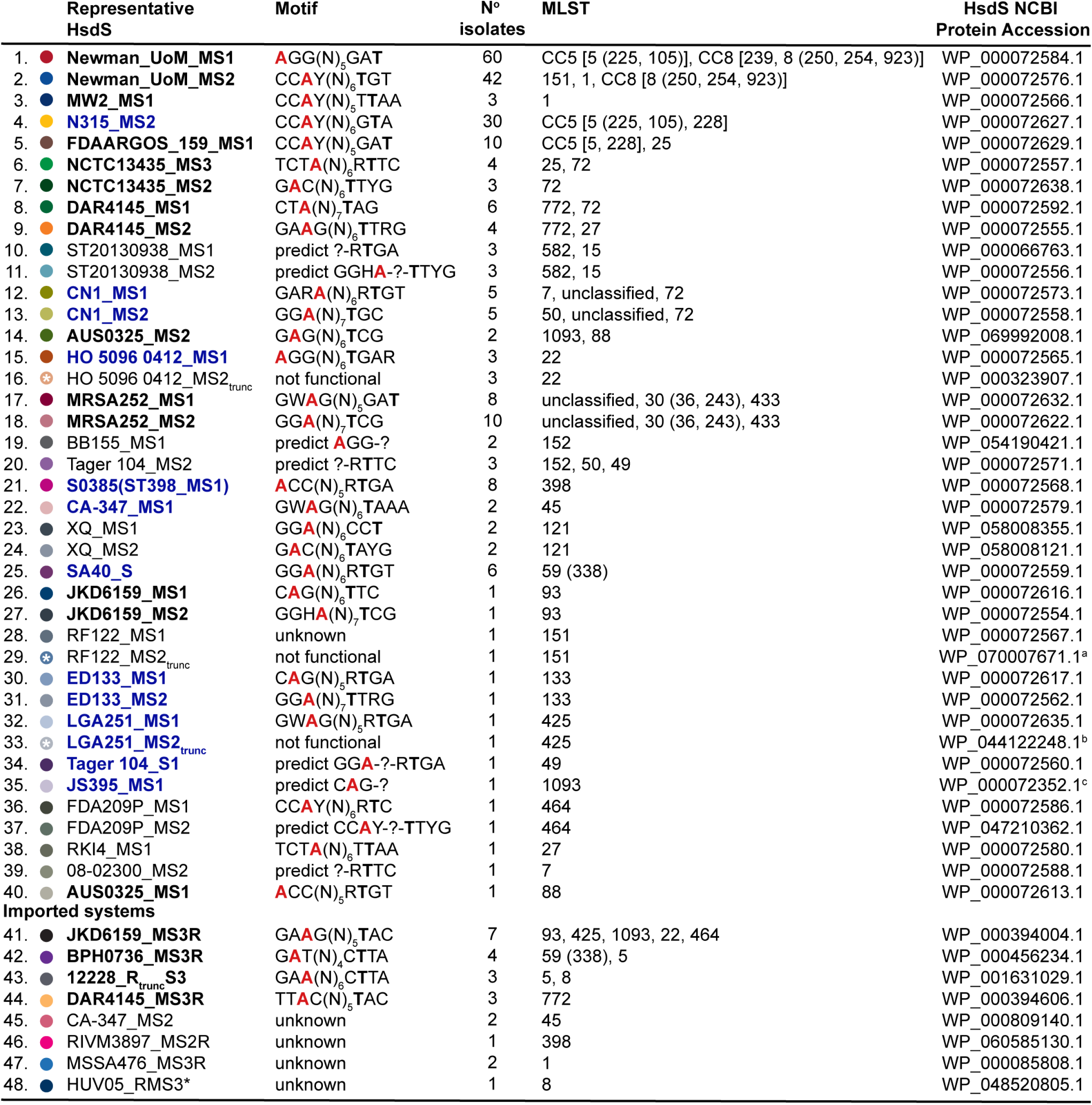
Diversity of S. aureus type I restriction modification system methylation profiles. Isolate HsdS motifs were collated from publications by Monk *et al.*, 2015 (9), Cooper *et al.*, 2017 (18) and the REBASE database (33). HsdS names in bold black font have motifs determined by PacBio sequencing of the isolate after which the representative HsdS was named. HsdS names in bold blue font have motifs determined by DNA cleavage with purified restriction enzyme. Multi-locus sequence types (MLSTs) in which each HsdS are found are listed according to the order they appear in Figure 2 phylogeny (top to bottom); STs within the same clonal complex (CC) are listed within square brackets; STs within parentheses are single locus variants of the ST group they are listed after. trunc = truncated; **A** (red) = methylated adenine residue; **T =** complementary partner to methylated adenine residue. ^a^ Truncation at amino acid 203. ^b^ Truncation at amino acid 249. ^c^ Truncation at amino acid 8. *HUV05_RMS3 is carried on a plasmid, not integrated in the chromosome. Full amino acid translations of all 48 HsdS variants are accessible at Figshare <u>DOI: 10.26188/5cb01fc089ab2</u>.

### *S. epidermidis* type I RM systems are carried on mobile genetic elements

Seven complete *S. epidermidis* reference genomes were publicly available at the beginning of this study (Table S1). Of these only BPH0662 (12) and RP62a had characterised type I RM system motifs. However the RP62a methylome was determined independently (11) of the finished genome (14). The methylomes of *S. epidermidis* isolates 1457 (15) and 14.1.R1 (16) confirmed that neither possessed a functional type I RM system, consistent with the absence of *hsdM* genes. To improve understanding of the type I RM systems in *S. epidermidis* we conducted PacBio SMRT sequencing and established complete genomes and adenine methylomes for six additional *S. epidermidis* strains from ST2, ST5, ST59 and ST358 and we re-sequenced RP62a (ST10) (See Table S3 for metadata).

The typical chromosomal arrangement of the type I RM system in *S. epidermidis* is shown for the BPH0736 (ST2) (Figure 1B). Unlike *S. aureus* (Figure 1A), in *S. epidermidis* type I RM systems are arranged as a complete three gene operon in either an *hsdRMS* or *hsdMSR* organisation, unless interrupted (Figure S1). Analyses of the 11 closed *S. epidermidis* genomes containing type I RM systems demonstrated their co-occurrence with cassette chromosome recombinase (*ccr*) genes (with or without the presence of *mec*). The 16 type I RM systems present within these 11 genomes were located a mean distance of 11.5 kb (minimum 2.3 kb, maximum 51.0 kb) from the nearest *ccr* (Figure S1A). Similarly, in the 22 *S. aureus* genomes with a SCC-associated type I RM system, the mean distance between *hsdRMS/MSR* and the nearest *ccr* was 6.9 kb (minimum 1.6 kb, maximum 20.9 kb) (Figure S1B).

Cassette chromosome recombinases typically integrate at *orfX* (corresponding to the last 15 nucleotides of the rRNA large subunit methyltransferase (17)) that is located at 31.6 kb in the *S. epidermidis* chromosome (33.3 kb in *S. aureus*). This is the start of a highly plastic region of the chromosome, in which multiple antibiotic resistance genes, drug transporters and insertion sequence (IS) elements have accumulated (12) (Figure S1). All 11 *S. epidermidis* and 22 *S. aureus* with *ccr-* associated type I RM systems were integrated at *orfX* (Figure S1). For type I RM system variants present in multiple isolates, conservation of genes surrounding the system and *ccr* was observed, consistent with the mobilisation of an entire element (Figure S1). Preserved cassette structure between isolates and both species (Figure S1B) led us to hypothesise that the movement of type I RM systems in *S. epidermidis* is mediated by *ccr*, enabling mobilisation on SCC elements between strains and to other staphylococcal species. Localisation in this region of the genome also predisposes *S. epidermidis* type I RM systems to disruption, potentially rendering variants restriction deficient. This is seen with interruption of *hsdR* by IS elements in BPH0736 (Figure 1B).

### *S. epidermidis* type I RM systems are not strictly conserved within lineages

To expand the *S. epidermidis* dataset we added short read data for 234 publicly available *S. epidermidis* genomes to the 13 finished genomes (Table S1). Unlike *S. aureus*, variability was noted in both the HsdR and HsdM subunits for *S. epidermidis*. Across the 247 genomes, 183 intact HsdR were identified, with five major variants (<90% amino acid pairwise identity threshold) (Table S2). The two variants of HsdR in strain BPH0662 shared only 22% amino acid identity. Similarly, 178 complete HsdM genes were identified with six major variants (Table 3; Table S2). The amino acid sequences of these six variants were markedly divergent. The two variants of HsdM present in BPH0662 shared only 31% amino acid identity.

**Table 3.**
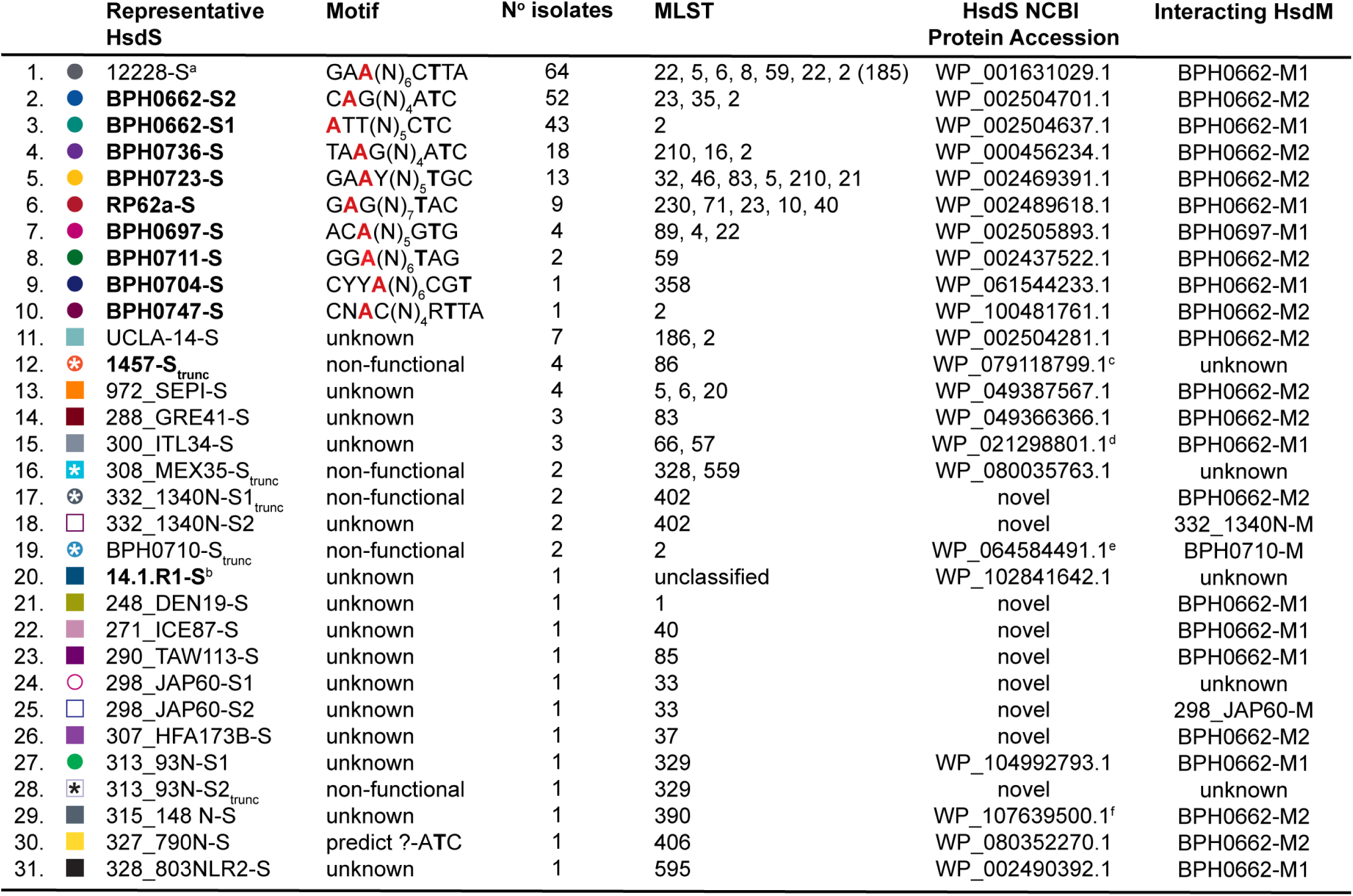
Diversity of *S. epidermidis* type I restriction modification methylation profiles. Isolate HsdS motifs were collated from methylomes newly characterised in this study and publications by Lee *et al.,* 2016 (12) and Costa *et al.,* 2017 (11). HsdS names in bold black font have motifs determined by PacBio sequencing of the isolate after which the representative HsdS was named. Multilocus sequence types (MLSTs) in which each HsdS are found are listed according to the order they appear in Figure 3 phylogeny (clockwise); ST185 is a single locus variant of ST2. trunc = truncated; **A** (red) = methylated adenine residue; **T =** complementary partner to methylated adenine residue. ^a^ ATCC 12228 type I RM system is non-functional with truncated *hsdR*, complete *hsdS* and no *hsdM*; all 64 isolates possessed the same incomplete type I RM system; Motif identified based on methylome for NIH4008 due to presence of HsdM capable of interacting with 12228 HsdS. ^b^ 14.1.R1 type I RM system is non-functional with truncated *hsdR*, complete *hsdS* and no *hsdM.* ^c^ L1M substitution. ^d^ 1^st^ 81 amino acids truncated. ^e^ S295P substitution. ^f^ 11 amino acid substitutions (K26E, I 56V, E59K, E171K, K174R, K175T, E178A, I193V, D201N, Y386F, V434I). Amino acid translations of all 31 HsdS variants (<u>DOI: 10.26188/5cb01e8f23305</u>) and their interacting HsdM (<u>DOI: 10.26188/5cb019b506f53</u>) are accessible through Figshare.

A maximum likelihood phylogeny for the 247 *S. epidermidis* genomes, derived from 83,210 core-SNPs, sampled from 72 STs was established and the 31 different *S. epidermidis* HsdS subunits identified were overlaid (Figure 3). Where known, their associated TRMs are shown in Table 3 with NCBI protein accession numbers (Table S2). The distribution of *S. epidermidis* HsdS within the population differed markedly from that observed within *S. aureus*, with no strict concordance to lineage specificity. For example, the HsdS from BPH0723 (BPH0723-S; Table 3) was present in 13 isolates from five STs (ST5, ST21, ST46, ST210, and one unclassified), while BPH0662-S2 was identified in 52 isolates from three STs (ST2, ST23 and ST35). Although a high proportion of ST2 isolates shared the same predicted methylome (Figure 3), the majority of these were known to be clones of internationally disseminated, multidrug resistant strain BPH0662 (5). However, even within this highly clonal group (n = 36), some predicted methylation variation existed. For example, BPH0662-S2 was absent from two isolates, six isolates (including BPH0662) had a truncation in the 12228-S orphan system while two isolates were missing the 12228-S orphan system completely. Furthermore, within ST2, seven different variants of HsdS were identified in 11 arrangements, including the absence of any type I RM system (Figure 3). Of the 247 *S. epidermidis* genomes analysed, 38% did not contain any *hsdS* alleles and were predicted to be restriction deficient.

**Figure 3.**
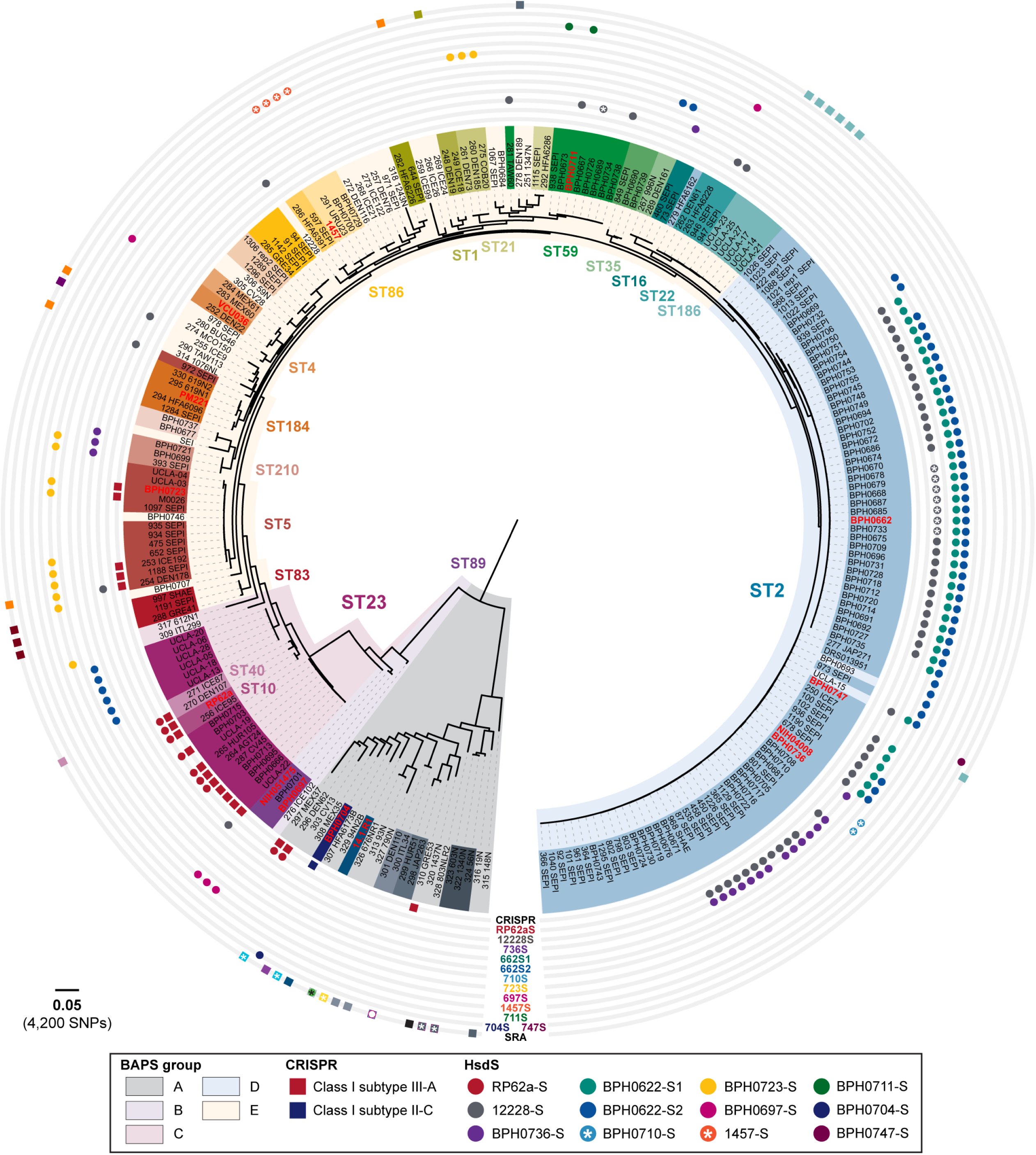
*S. epidermidis* type I restriction modification systems are not conserved within lineages. Maximum-likelihood, core-SNP based phylogeny for 247 *S. epidermidis* genomes: seven newly closed reference genomes; six existing reference genomes; 156 genomes curated from NCBI sequence read archive (SRA); 75 isolates from Lee *et al*. (5); and the three draft genomes with methylation data (11). BPH0736 was used as the reference genome for analyses. Overlaid are the results of in silico multi-locus sequence type (MLST), bayesian analysis of population structure (BAPS), presence of CRISPR-Cas systems and type I restriction modification system HsdS variants. Bold red font indicates isolates with characterised methylomes. Isolates were from 70 recognised and two unclassified MLST groups. Boxes around strain names are coloured according ST type; when background colour is same as BAPS group, indicates an ST represented by a single isolate. *Truncated HsdS subunit. Scale bar indicates number of nucleotide substitutions per site (bold) with an approximation of SNP rate (in parentheses).

Our genome sequencing of RP62a_UoM indicated the presence of a single type I RM system with a G**A**GN_7_TAC TRM (Table 3). Although this motif was consistent with that previously reported by Costa *et al*., the three additional motifs previously described (lacking apparent associated genes) were not detected by our methylome analysis; the low complexity of the motifs (e.g. GGBNNH) and low frequency of detected methylation (12-29%) (11) suggest these may have been artefacts rather than true motifs. Similarly, the three additional low complexity and frequency motifs reported for VCU036 (11) were probable artefacts. Although the ST type was not specified, VCU036 that shared the same methylome as ST89 isolate NIH051475 was reported as CC89 by Costa *et al.* leading to the conclusion that *S. epidermidis* type I RM systems follows *S. aureus-*like lineage specificity (11). We performed *in silco* MLST by two independent methods and determined VCU036 to belong to ST4. Furthermore, our analysis of the 247 *S. epidermidis* genomes demonstrated VCU036 to be phylogenetically distinct from ST89 (Figure 3).

Overall, our analyses demonstrated that contrary to current assumptions (11) the type I RM systems of *S. epidermidis* do not adhere to the lineage specific distribution observed in *S. aureus*. These differences are attributable to the arrangement of *S. epidermidis* type I RM systems as complete three gene operons that reside within a highly mobile region of the chromosome, the movement of which we hypothesise to be mediated by *ccr*.

### Recombinant target recognition domains generate HsdS variants with low conservation of amino acid identity

The structure of a typical type I RM system HsdS allele is shown in Figure S2A, composed of two highly variable target recognition domains (TRDs) flanked and separated by conserved regions (CRs) that collectively determine the TRM to be methylated by HsdM. Recombinant pairings of TRDs result in different variants of HsdS (13, 18). Alignments for the range of *S. aureus* and *S. epidermidis* HsdS identified in this study are shown in Figures S2B and S3 respectively. Within our *S. aureus* and *S. epidermidis* collections, 77 variants of HsdS that shared only 24% pairwise identity were identified. This low conservation poses a potential challenge to the high throughput bioinformatic screening for HsdS variants within genomic datasets. However, using the HsdS from ATCC 12228 as the reference translation with our described method, we were able to detect the partial if not complete presence of all HsdS variants in both species. Of note, 12228-S was the only HsdS variant found within both species that clustered with the majority of *S. aureus* variants. In comparison, RP62a-S only captured 18 of the 31 *S. epidermidis* HsdS variants and fragments of under half the *S. aureus* HsdS variants.

### *S. epidermidis* HsdS variants will only interact as part of a specific complex

The arrangement of some *S. epidermidis* type I RM systems, with the presence of a truncated *hsdS* between complete *hsdR* and *hsdMS* genes, suggested that recombination of component genes occurs (e.g. SepiBPH0662I, SepiRP62aI and SepiBPH0704I; Figure S1). Analyses of the 247 genomes indicated that each variant of *hsdS* in *S. epidermidis* is always associated with a specific *hsdR* and *hsdM* gene, with a conserved gene arrangement (unless interrupted), frequently with the same surrounding genes in association with *ccr* (Figure S1A). These observations support our hypothesis for a role for SCC elements in the mobilisation of *S. epidermidis* type I RM systems.

The presence of an orphan *hsdS* without a partner *hsdM* gene in *S. epidermidis* introduces additional complexity to the prediction of type I RM system functionality. This was demonstrated by 12228-S, the most prevalent HsdS within the dataset, that is found in 64 *S. epidermidis* (Table 3, Figure 3) and three *S. aureus* (Table 2, Figure 2) isolates. All examples of this *hsdS* variant followed a truncated *hsdR*, without an *hsdM* gene. We determined that 12228-S is only expressed if its specific interacting variant of *hsdM* (BPH0662-M1; WP_002504638.1) is also present (see Table S4 for full explanation). In contrast, conservation of a single variant of *hsdM* present twice within the same *S. aureus* genome provides redundancy for the expression of type I RM methylation. This is consistent with previous findings where the product of a single copy of the conserved *S. aureus hsdM* allele could functionally interact with both CC8 HsdS when heterologously expressed in *E. coli* (9).

### Plasmid artificial modification to overcome the type I RM systems in *S. epidermidis* provides equivalent transformation efficiency as deletion of functional type I systems

To determine the restriction barrier posed by type I RM systems in *S. epidermidis* and assess the efficiency of PAM as a means of bypassing restriction barriers (Figure 4), Δ*hsdS* mutants and *E. coli* hosts for PAM were constructed for *S. epidermidis* isolates BPH0662, RP62a and BPH0736. Two different plasmids (pRAB11 (19) and pIMAY (8)) were used in transformation experiments, as each carried a different number of TRMs recognised by the type I RM systems present in each isolate (Figure 4D). Clinical ST2 isolate, BPH0662-WT, was found to have an intractable restriction barrier unless both functional type I RM systems were overcome through complete bypass with PAM in an *E. coli* host (Ec_Se662I-II), by deletion of both complete *hsdS* genes (BPH0662Δ*hsdSI*Δ*hsdSII*), or a combination of both approaches (plasmid from Ec_Se662I transferred into BPH0662Δ*hsdSII* or plasmid from Ec_Se662II transferred into BPH0662Δ*hsdSI*) (Figure 4A).

**Figure 4.**
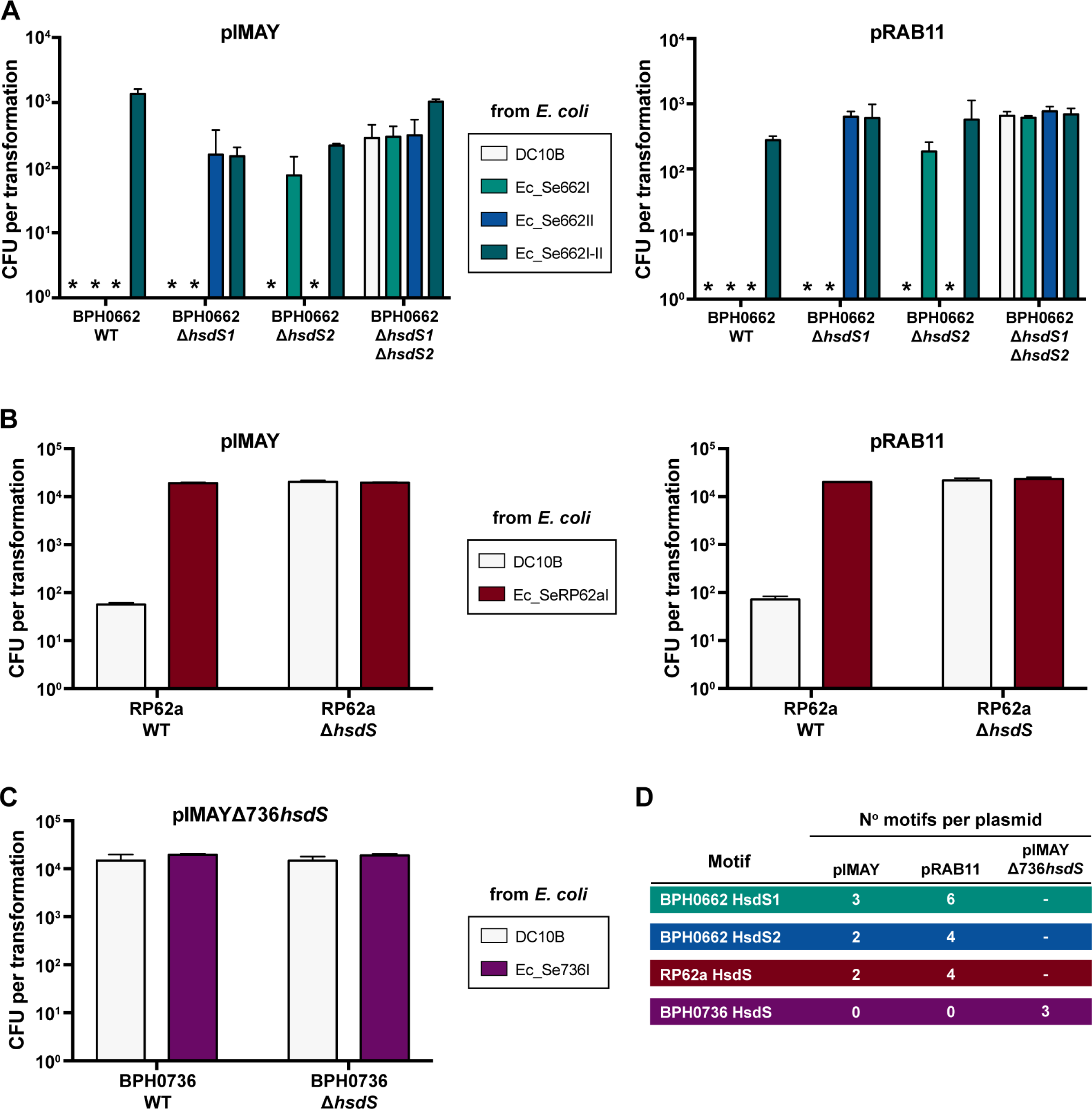
Plasmid artificial modification to overcome the type I RM systems in *S. epidermidis*. Biological triplicate data for 5 ug of plasmid passaged through DC10B *E. coli* compared to the relevant *E. coli* PAM construct, transformed into *S. epidermidis* wild type (WT) and Δ*hsdS* mutant strains. Error bars represent mean ± standard deviation of three independent experiments. CFU = colony forming units. *No transformants. **A.** Transformation of BPH0622-WT, BPH0662Δ*hsdS1*, BPH0662Δ*hsdS2* and BPH0662Δ*hsdS1*Δ*hsdS2* with plasmid pIMAY or pRAB11 isolated from DC10B and strain specific *E. coli* Ec_Se662I (expressing BPH0662*hsdMS1*), Ec_Se662II (expressing BPH0662*hsdMS2*) and Ec_Se662I-II (expressing both BPH0662*hsdMS1* and BPH0662*hsdMS2*). **B.** Transformation of RP62a-WT and RP62aΔ*hsdS* with plasmid pIMAY or pRAB11 isolated from DC10B and strain specific *E. coli* Ec_SeRP62aI (expressing RP62a*hsdMS*). **C.** Transformation of BPH0736-WT and BPH0736Δ*hsdS* with plasmid pIMAYΔ736*hsdS* isolated from DC10B and strain specific *E. coli* Ec_Se736I (expressing BPH0736*hsdMS*). Note, pIMAYΔ736*hsdS* was used as neither pIMAY nor pRAB11 possessed any TRM. **D.** Number of *S. epidermidis* strain specific HsdS TRM present on each plasmid.

Using our protocol, the type I restriction barrier in RP62a-WT was found to be incomplete. Low numbers of transformants (10^1^ CFU/ml) were obtained with plasmid DNA isolated from DC10B, indicating that bypassing the type IV restriction barrier alone was sufficient to allow genetic manipulation of this strain (Figure 4B) as previously demonstrated (8). Complete bypass of the single type I RM system in this isolate with *E. coli* host Ec_SeRP62aI significantly improved transformation efficiency to 10^4^ CFU/ml, which was equivalent to complete absence of a functional type I RM system as determined with the RP62aΔ*hsdS* mutant (Figure 4B). In contrast, when expressing the RP62a *hsdMS* genes from a plasmid in DC10B, Costa *et al*, were unable to completely bypass the type I RM barrier. This discrepancy was attributed to additional RM systems with low frequency methylation (11). However, our results showed that only one type I RM system is present in RP62a, suggesting that the heterologous expression of type I RM systems on a plasmid in DC10B rather than from a single copy of the genes integrated into the chromosome may be suboptimal. Previously, we found that plasmid-based expression of *hsdMS* was both unstable and cells were unable to tolerate high level expression required for complete methylation of the target DNA (Monk et al 2015).

Clustered, regularly interspaced, short palindromic repeat (CRISPR) loci confer sequence directed immunity against phages and other foreign DNA, and are another recognised barrier to horizontal gene transfer in *S. epidermidis* (20). Our analysis of the CRISPR spacers for RP62a (Table S3D) did not demonstrate the presence any targets on pSK236 (5.6 kb) used by Costa *et al*., or on pRAB11 (6.4 kb) or pIMAY (5.7 kb) used in this study, that would account for their lower transformation efficiency (10^2^ CFU/ml per 5 µg plasmid DNA) (11) compared to our protocol (10^4^ CFU/ml per 5 µg plasmid DNA for both pRAB11 and pIMAY).

Isolate BPH0736-WT was predicted to be naturally restriction deficient due to the interruption of *hsdR* by IS elements (Figure 1B) but was demonstrated to retain functional methylation conferred by an intact *hsdMS* system. Due to the complex and infrequently occurring TRM dictated by the single *hsdS* (Table 3), neither pIMAY nor pRAB11 had any BPH0736-S TRMs present. Therefore, pIMAY bearing the Δ736*hsdS* insert (pIMAYΔ736*hsdS*) was used as this contained three TRMs (Figure 4D). BPH0736-WT was functionally confirmed to be restriction deficient with the same transformation efficiency of 10^4^ CFU/ml demonstrated for both BPH0736-WT and BPH0736Δ*hsdS* using plasmid isolated from non-specific *E. coli* host DC10B and PAM tailored mutant Ec_Se736I (Figure 4C). Further supporting our bioinformatic predictions, like BPH0736, ATCC 12228 (truncated *hsdR* and no *hsdM*; ST8) and BPH0710 (truncation at amino acid 81of HsdS; ST2) were both predicted to have no functional restriction barrier. Similar to BPH0736, both these strains were transformable in the order of 10^4^ CFU/ml with plasmid isolated from DC10B, suggesting that this was the maximum transformation efficiency expected for our protocol. Clinical ST2 strain, BPH0676, was also predicted to have no restriction barrier with complete absence of a type I RM system, however similar to BPH0662 the maximum transformation efficiency achieved was only 10^3^ CFU/ml suggesting inherent strain dependent factors other than type I RM systems impacted on transformation of these isolates e.g. cell wall thickness (21).

Although the above data demonstrates PAM to be an efficient method to overcome the type I restriction barrier of *S. epidermidis*, we observed potential instability with the integration of multiple *S. epidermidis hsdMS* of particular TRMs in a DC10B *E. coli* background. With serial passage of Ec_Se662I-II the transformational efficiency of plasmid isolated from this *E. coli* host into BPH0662 declined from 10^3^ to 10^1^ CFU/ml despite all other experimental parameters remaining the same. This was not observed for any of the *E. coli* PAM mutants expressing single *hsdMS*, including Ec_Se662I and Ec_Se662II, that maintained high-level methylation (89.65 – 99.90%) of motifs within the genome (12) (Table S3). Illumina sequencing of the Ec_Se662I-II genome confirmed integration of both *hsdMS* at the expected chromosomal sites but loss of approximately half the coding sequence of both *hsdS* genes for the majority of the population sequenced. This instability was hypothesised to be due to the burden of excessive DNA methylation (10,930 sites from heterologous expression of two BPH0662 *S. epidermidis* type I RM systems in addition to 38,592 sites of endogenous *E. coli dam* methylation) that may interfere with normal cellular function, rendering expression toxic in *E. coli*. The same likely accounts for the poor transformation efficiency reported when using PAM for NIH4008 (100-fold lower than that observed for isolates with only a single type I RM system) by Costa *et al*. (11). NIH4008 possesses the same type I RM systems as BPH0662, without the truncation of the orphan *hsdS* (Figure 3). Furthermore, although stable chromosomal integration of three *S. aureus hsdMS* systems in *E. coli* DC10B (IM93B) has been described by Monk *et al*., decreased efficiency of methylation was observed with only 10,135 of 14,602 total TRM sites demonstrating detectable methylation (9).

Collectively, our current and previous (9, 12) data suggests that DC10B *E. coli* is unlikely to consistently maintain the heterologous expression of staphylococcal type I RM systems in the setting of high frequency methylation (>10,000 sites). This limitation should not impact plasmid transformation for mutant creation by allelic exchange, which theoretically requires only a single transformant. However, should high efficiency transformation be sought (e.g. direct transposon mutant library selection), then suitable strains can be predicted using genomic data to identify restriction deficient isolates, such as our newly described reference isolate BPH0736, a clinically significant, ST2 isolate. Clinical metadata, genome characteristics, CRISPR spacers (when present), *in silico* resistome, Vitek 2 antibiogram for clinically relevant antibiotics and common plasmid selection markers for the seven new reference isolates and BPH0662 are shown in Table S3. Metadata and sequencing accession for mutant isolates is listed in Table S5.

### Phage transduction of plasmid is subject to type I restriction

Phage transduction is an alternate method for the genetic manipulation of *S. epidermidis*. Particularly, *S. aureus* ST395 lineage specific Φ187, that shares wall teichoic acid (WTA) receptors with *S. epidermidis* (22, 23). Dependent upon the incidental packaging of plasmid introduced into restriction deficient intermediary host, *S. aureus* PS187Δ*hsdR*Δ*sauPSI*, with Φ187 phage machinery (24), the method can be used to transduce a number of CoNS but is not universally applicable to all *S. epidermidis* isolates (23). The observed ability of ST395 *S. aureus* to exchange DNA with some CoNS led Winstel *et al.* to conclude that overlap may exist between the DNA methylation of ST395 *S. aureus* and CoNS that share the same WTA receptors (22, 23). Phage Φ187 transduction experiments performed using our WT isolates and Δ*hsdS* mutants for BPH0662, RP62a and BPH0736 are shown in Figure 5 examining the transfer of pRAB11. These experiments demonstrate that even if successfully transduced into a *S. epidermidis* isolate, plasmids are still subject to degradation by type I RM systems if they bear a recognised TRM. However, in BPH0736 (absent type I restriction) or mutant strains in which systems have been rendered inactive, transduced plasmid remains viable. The methylome for ST395 *S. aureus* has not been characterised, however the draft genome sequence for PS187 (GCA_000452885.1) indicates that both type I RM systems in this isolate are identical to those in *S. aureus* isolate JS395 (ST1093, belonging to CC395 (25)). We predicted the methylome of the isolate to include G**A**GN_6_TCG (same as AUS0325-MS2) and another unknown TRM (Figure 2, Table 2). The results of our experiments and analyses of the diversity of *S. epidermidis* type I RM systems suggest that successful phage transduction of some *S. epidermidis* isolates with Φ187 is more likely related to the absence of a functional system, rather than the presence of a shared methylome with ST395 *S. aureus*. This is further supported by experiments performed by Winstel *et al.*, (23) in which Φ187 could only transduce pTX15 (26) into RP62a, but not pKOR1 (27). Based on our characterised RP62a TRM, we determined that pTX15 possesses no RP62a motifs, while both pRAB11 and pKOR1 each bear four motifs explaining why neither plasmid is transducible into RP62a-WT.

**Figure 5.**
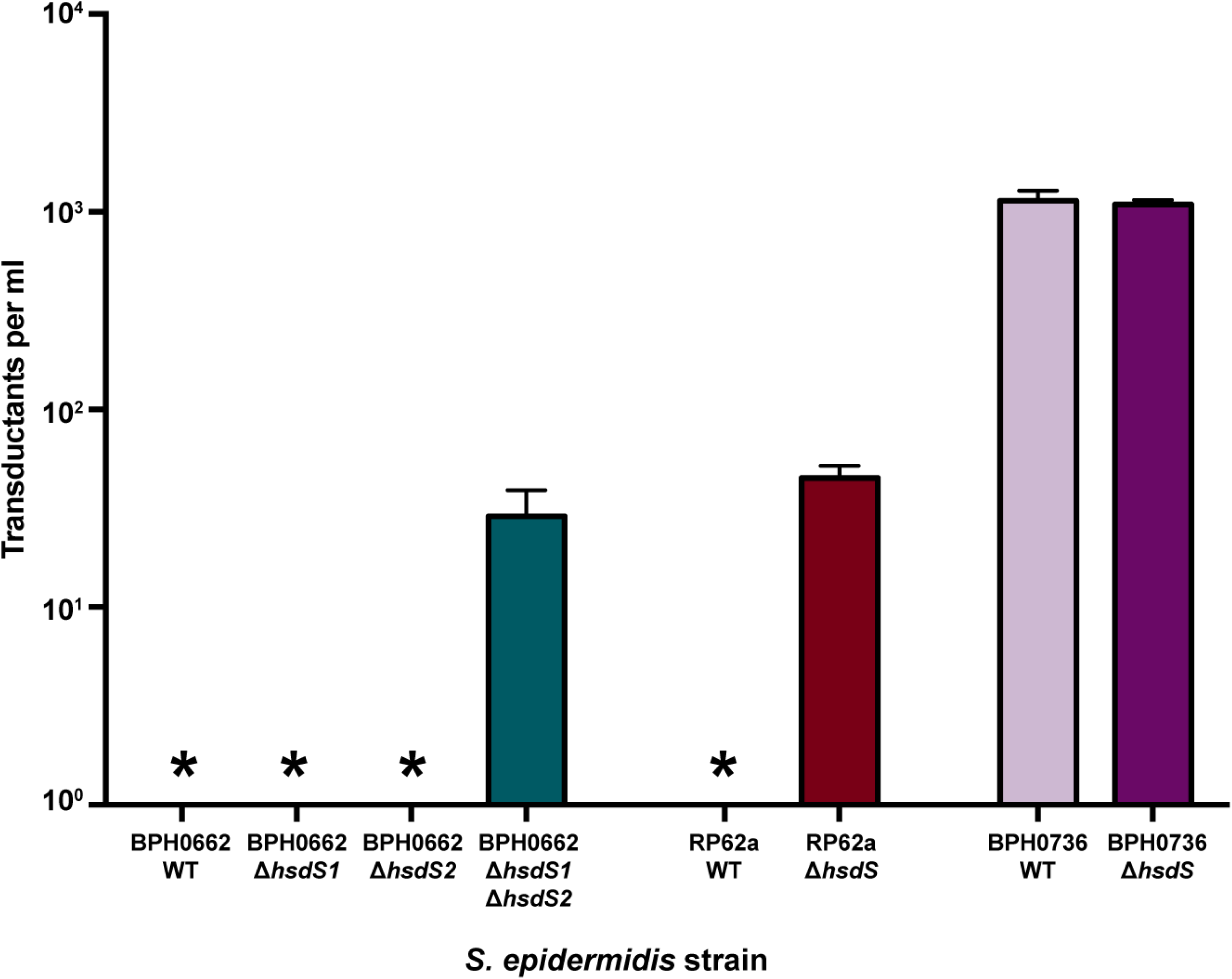
*S. epidermidis* phage transduction is subject to type I restriction. Biological triplicate data for phage transduction of Φ187-pRAB11 lysate transduced into *S. epidermidis* wild type (WT) and *hsdS* mutant strains. Error bars represent mean ± standard deviation of three independent experiments. *No transformants.

Temperature sensitive plasmid, pIMAY, is frequently used for allelic exchange in staphylococci due to the presence of inducible *secY* antisense counter selection, and the lower likelihood of unintended mutations (such as occur with pKOR1) as integrants are selected at 37°C instead of 43°C (28). However, we found that Φ187 was not capable of transducing pIMAY into any of the tested strains, including the Δ*hsdS* mutants and naturally restriction deficient BPH0736. We hypothesised this was due to the low copy number of pIMAY in staphylococci, resulting in low levels of incidental packaging of the plasmid within Φ187, as compared to high copy number plasmid pRAB11. Other limitations to Φ187 transduction include a recommendation to use plasmids <10 kb (24), that should not have impacted on pIMAY (5.7 kb), which is smaller than pRAB11 (6.4 kb). Although a simplified harvesting and infection protocol was used compared to that described by Winstel *et al.* (24), in restriction deficient *S. epidermidis* strain BPH0736 we achieved an efficiency of 10^4^ transductants per ml, equivalent to their anticipated results of 10^1^-10^4^ (24). Of note, the efficiency of the Δ*hsdS* mutants, BPH0662Δ*hsdSI*Δ*hsdSII* and RP62aΔ*hsdS* were both two-log lower compared to BPH0736 (Figure 5), further supporting the presence of strain dependent factors beyond the barriers posed by type I RM systems and WTA in these backgrounds.

## Conclusions

Our results demonstrate marked differences between the type I RM systems in *S. aureus* and *S. epidermidis* that had hitherto been assumed to be same (11). These differences are predominantly attributable to the arrangement and genome location of the *S. epidermidis* type I system as a complete three-gene operon, we hypothesise to be mobilised by *ccr*. Localisation of the operon in a highly plastic region of the chromosome increases the likelihood of horizontal transfer of these complete systems between *S. epidermidis* as well as to other staphylococci. This results in a lack of lineage specificity and higher probability of spontaneous interruption of component genes. This is in contrast to *S. aureus* where the type I systems are typically arranged as one *hsdR* and two *hsdMS* genes located distant from one another in stable regions of the chromosome. The diversity of *S. epidermidis* type I RM systems that do not strictly adhere to ST/CC groupings indicates that genetic manipulation of *S. epidermidis* requires tailoring to each isolate of interest. Attempting transformation without genomic analysis of the methylome could be successful, as our analyses found that 38% of *S. epidermidis* strains did not possess a type I RM system, and not all systems pose an intractable barrier (e.g. RP62a). However, some isolates such as the internationally disseminated, near pan-drug resistant, clone BPH0662 have complex and absolute type I restriction barriers.

We have demonstrated that PAM using a DC10B *E. coli* host is a simple and effective means to bypass the type I RM barrier in *S. epidermidis*, with a plasmid transfer efficiency equivalent to a complete absence of type I RM systems. The decreasing cost and ready availability of whole genome sequencing has made the sequencing of isolates planned for mutagenesis and their mutant derivatives commonplace, and a practice that is recommended to ensure the absence of acquired secondary mutations (29). If the genome sequence of an isolate is known, its methylome and ability to be transformed can be predicted as follows: 1. Does the isolate possess an intact type I RM system? If not type I methylation will not be expressed and the isolate should be inherently transformable. 2. Each complete type I RM system within a genome should be functional. For an HsdS protein with known TRMs, presence of the TRMs on a vector will likely prevent transformation. 3. Orphaned, complete *hsdS* genes in the absence of an adjacent *hsdM* may be expressed if their associated *hsdM* allele is present elsewhere in the genome. In view of the above, when designing an *E. coli* PAM host, we recommend including all complete *hsdMS* and any complete orphan *hsdS* genes from the *S. epidermidis* strain to be manipulated, to ensure complete recapitulation of the endogenous type I methylome.

The 247 genomes we analysed are by no means an exhaustive representation of all *S. epidermidis* and additional examples of type I RM systems will undoubtedly be catalogued as further sequencing of this organism is performed. However, this genomic sampling together with our functional data was sufficient to draw the above conclusions. In view of the identified complexities associated with the genetic manipulation of *S. epidermidis*, the reference isolate BPH0736 should prove particularly useful. A clinical ST2 isolate representative of international circulating clones (5), BPH0736 is naturally transformable due to the spontaneous interruption of *hsdR*, rendering it highly amenable to both transformation and phage transduction, making it an ideal strain for future molecular studies.

## Materials and Methods

### Media and reagents

Bacterial strains, plasmids and oligonucleotides used in this study are listed in Table S6. *S. epidermidis* were routinely cultured at 37°C in brain heart infusion broth (BHI)(Difco). See Supplementary Methods for detailed description of culture media, antibiotics and enzymes.

### Genome sequencing and analysis

Refer to Supplementary Methods.

### Electroporation

Early stationary phase cultures (8 h) of *S. epidermidis* grown in 10 ml of B Media (BM) were added to 90 ml of fresh, prewarmed BM. Cultures were reincubated to an OD_600_ between 0.8 - 0.9 and chilled in an ice slurry for 10 min. Cells were harvested at 3,900 *xg* for 5 min at 4°C in a swinging bucket rotor and the cell pellet resuspended in 100 ml of autoclaved, ice-cold water. Centrifugation was repeated, and the pellet resuspended in 50 ml of autoclaved ice-cold water. Cells were centrifuged and successively resuspended in 20 ml, 10 ml then 250 μl of autoclaved ice-cold 10% (weight/volume) glycerol. Equal aliquots (50 μl) were frozen at −80°C. Prior to electroporation, cells were thawed on ice for 5 min, then at room temperature for 5 min. Following centrifugation at 5,000 *xg* for 1 min, cells were resuspended in 50 μl of 10% glycerol with 500 mM sucrose (filter sterilised). Pellet paint (Novagen) precipitated plasmid DNA was added to the cells, then cells were transferred into a 1 mm electroporation cuvette (Bio-Rad) and pulsed at 21 kV/cm, 100 Ω, 25 μF at room temperature. Routinely, 5 μg of plasmid DNA was used, with concentration determined by fluorometric assay (Qubit 2.0; Life Technologies). Cells were incubated in 1 ml of BHI supplemented with 500 mM sucrose (filter sterilised) at 28°C for 2 h prior to plating on BHIA containing chloramphenicol 10 μg/ml.

### Construction of Ec_Se736I and Ec_SeRP62aI *E. coli* hosts

*E. coli* mutants expressing the relevant *S. epidermidis* type I RM systems in a DC10B background were created using the primers listed in Table S6 as previously described (8, 9, 12). Detailed methodology in Supplementary Methods.

### Construction of *S. epidermidis* Δ*hsdS* mutants

The pIMAY(Δ*hsdS*) vectors were constructed using amplified by overlap extension PCR (30) with the A/B/C/D primer sets specified for each strain in Table S6; cloning into the pIMAY vector backbone; subsequent cloning of the insert into the vector; mutant selection and screening were conducted as previously described (5). Detailed methodology in Supplementary Methods.

### Harvesting Φ187 + pRAB11/pIMAY lysate from *S. aureus* PS187Δ*hsdR*Δ*sauPSI*

Φ187 containing pRAB11/pIMAY was harvested from *S. aureus* PS187Δ*hsdR*Δ*sauPSI* using a protocol adapted from Winstel (24). See Supplementary Methods for detailed methodology.

### Φ187 + pRAB11/pIMAY transduction of *S. epidermidis*

A phage transduction protocol was adapted from Foster (31). Detailed methodology in Supplementary Methods.

### Accession numbers

The datasets supporting the results of this article are available from NCBI under BioProject No. PRJNA532483 (sequencing and closed genome assemblies) and Figshare (*<u>S. aureus</u>* <u>HsdS</u>; *<u>S. epidermidis</u>* <u>HsdS</u>; *<u>S. epidermidis</u>* <u>HsdM</u>; *<u>S. epidermidis</u>* <u>HsdR</u>).

### Funding information

This project was supported by the Royal Australasian College of Physicians, Basser Research Entry Scholarship/Australian Government Research Training Program Scholarship to JYHL; National Health and Medical Research Council of Australia (NHMRC) Senior Research Fellowship to TPS (GNT1105525); NHMRC Practitioner Fellowship to BPH (GNT1105905); and NHMRC Project Grant (GNT1066791).

## Acknowledgments

We thank Volker Winstel for supply of Φ187 and *S. aureus* strain PS187Δ*hsdR*Δ*sauPSI*. We also acknowledge Bernhard Krismer for providing the assembled DNA sequence of pTX15.

## Author contributions

Project conceived by JYHL, TPS, BPH and IRM. Experimental work performed by JYHL and IRM, with PacBio sequencing performed by GPC and SJP. Bioinformatic analysis performed by JYHL with assistance from TS, RG and AGdS. JYHL, TJF, BPH, TPS and IRM drafted the manuscript; all authors reviewed and contributed to the final manuscript.

---

**Figure S1. A hypothesised role for cassette chromosome recombinase (*ccr*) in the mobilisation of *S. epidermidis* and *S. aureus* type I restriction modification systems. A.** Complete *S. epidermidis* genomes with type I RM systems. **B.** *S. aureus* genomes with imported type I RM systems. Genomes are orientated forwards starting at *dnaA*. ^#^NCBI uploaded genome does not start at *dnaA*. *SEI not classifiable by existing MLST scheme.

**Figure S2. Alignment of *S. aureus* HsdS variants. A.** Structure of an HsdS allele with conserved regions (CRs) flanking two variable regions known as target recognition domains (TRD1 & TRD2). **B.** Each TRD typically specifies three to four defined base pairs including a methylated adenine residue (red **A**; **T** = complementary partner to methylated adenine residue); with a four to seven base pair non-specific spacer (N) between the two defined halves, collectively these TRDs determine the full target recognition motif (TRM) specified by an HsdS variant. HsdS names in bold black font have motifs determined by PacBio sequencing of the isolate after which the representative HsdS was named. HsdS names in bold blue font have motifs determined by DNA cleavage with purified restriction enzyme. Alignments of the identified variants of *S. aureus* HsdS are shown adjacent to their TRMs, each formed by a different TRD pairing. Scale above alignments indicates the position in the consensus alignment with mean pairwise identity at each site graphed (green = 100% identity; khaki = 30-100%; red <30%). Blue (TRD1) and red (TRD2) outlines highlight examples of TRDs that recur within the alignments and the TRM base pairs they define. Yellow boxes highlight alignments of HsdS imported into *S. aureus* on Staphylococcal cassette chromosome elements.

**Figure S3. Alignment of *S. epidermidis* HsdS variants.** HsdS names in bold black font have motifs determined by PacBio sequencing of the isolate after which the representative HsdS was named. Target recognition motifs (TRMs) (when known) and amino acid alignments are shown adjacent. Scale above alignments indicates the position in the consensus alignment with mean pairwise identity at each site graphed (green = 100% identity; khaki = 30-100%; red <30%). Red outline highlights an example of a target recognition domains that recurs within the alignments and the TRM base pairs they define. Yellow box highlights the 12228-S alignment, this HsdS variant was found to be present in both *S. aureus* and *S. epidermidis* and shared conserved regions with the *S. aureus* HsdS variants located within stable chromosomal islands (Figure S2).

**Table S1. Metadata associated with pre-existing S. aureus and S. epidermidis sequencing. A.** S. aureus reference genomes; **B.** S. epidermidis reference genomes; **C.** S. epidermidis SRA isolate metadata; **D.** Lee *et al*., 2018 isolate metadata; **E.** Costa *et al*., 2017 isolate metadata.

**Table S2. *S. aureus* and *S. epidermidis* type I restriction modification systems subunit protein accession numbers and references. A.** *S. aureus* HsdS; B. *S. aureus* HsdR and HsdM; C. *S. epidermidis* HsdS; D. *S. epidermidis* HsdR and HsdM.

**Table S3. S. epidermidis reference isolates metadata. A.** Clinical metadata; **B.** Closed genome statistics; **C.** PacBio methylation statistics; **D.** CRISPR; **E.** Vitek 2 susceptibilities; **F.** Resistome; **G.** Susceptibilities to commonly used plasmid selection markers.

Table S4. Determining the HsdM variant that interacts with 12228 HsdS.

**Table S5. S. epidermidis ΔhsdS and E. coli plasmid artificial modification mutant sequencing accession data.** A. PacBio sequencing; **B.** PacBio methylation; **C.** Illumina sequencing.

Table S6. Strains, plasmids & oligonucleotides used in this study.

